# DUnet: A deep learning guided protein-ligand binding pocket prediction

**DOI:** 10.1101/2022.08.11.503579

**Authors:** Xinglong Wang, Beichen Zhao, Penghui Yang, Yameng Tan, Ruyi Ma, Shengqi Rao, Jianhui Du, Jian Chen, Jingwen Zhou, Song Liu

## Abstract

Investigating protein-ligand binding sites is the key step in engineering protein/enzyme activity and selectivity. In this study, we developed a 3D convolutional neural network DUnet that derived from DenseNet and UNet for predicting the protein-ligand binding sites. To train DUnet, the features of protein 3D structure were extracted by describing the atomic physical characters, and the ligand binding sites were used as training labels. DUnet was trained using three dataset, the scPDB dataset (collecting of protein-ligand complexes from Protein Data Bank), scPDB and SC6K (collecting of protein-ligand complexes deposited after January 1st, 2018 from Protein Data Bank) datasets, and scPDB and its derived dataset by rotating the samples in the dataset. DUnet displayed better performance than the current state-of-art methods during the benchmark test using independent validation sets, and enlarging the training set contributed to better accuracy. We developed a small dataset contains commonly used industrial enzymes for testing DUnet and found that it was also accurate in predicting the substrate binding sites. We experimentally characterized the substrate binding sites of microbial transglutaminase according to the prediction and showed the significance of these sites. Finally, DUnet was used to predict the ligand binding sites of Swiss-Prot annotated proteins.

## Introduction

Structural-based protein design has achieved great success in protein/enzyme engineering [1]. The formation of the protein-ligand binding pocket critically affects the specificity and activity of proteins. Three conventional steps are used for engineering protein specificity and activity: (1) obtaining an accurate protein structure either by crystallization or structural modeling; (2) recognition of the protein-ligand binding sites; (3) evaluating the binding energy changes upon mutagenesis. The thriving of accurate protein modeling tools like Alphafold-2 [2] and RosettaFold [3], and mutation free energy calculation tools like Rosetta ddG [4] and FoldX [5] have significantly benefited the protein engineering workflow. Predicting protein-ligand binding sites is the pre-step of virtual mutagenesis. Accurate recognition of the key binding sites can assist the design of protein pockets, and provide target sites for virtual mutagenesis. Therefore, developing a tool for fast and accurate finding the protein-ligand binding sites is greatly needed.

Classical methods are mainly template-, geometric- or probe-based [6-10]. Through using the template-based method, the protein-ligand binding sites was obtained by aligning target structure to their identical protein homologs with known protein-ligand interactions in the databases [6]. The template-based method such as FINDSITE requires enriched databases which covered annotate protein-ligand binding knowledge to give instructions to new structures. Geometric-based method targets the cavities in the protein structure, and the possibility of each cavity becoming binding sites was ranked. Fpocket is a geometric-based protocol that ranks the pocket based on Voronoi tesselation and α-spheres [7]. However, such protocol scores the pocket corresponded to protein structural information; hence, accurate prediction of protein structures is highly needed. The probe-based protocol can also be recognized as binding energy-based protocol. In tools like FTSite [8], the cavities within the protein are accommodated with 16 virtual probes, and the binding sites are ranked based on the binding energy between probes and protein cavities. However, the limitation of using this method is that the actual binding pocket and ligand may not meet the lowest binding energy rule, and virtual probes cannot fully represent the diversified compounds that exist naturally.

Convolutional neural network (CNN) is an effective and robust model for resolving segmentation problems [11]. Recently, CNN has been implemented for classification purposes in biological fields, such as for segmenting 2D bacterial images [12], diagnosis of gastroscopic biopsy [13] or3-dimensional (3D) MRI images [14]. For segmenting protein and protein-ligand binding sites in a 3D image, the reported 3D-CNNs models including Kalasanty, Deeppocket, and PUResNet showed better performance than the traditional physics-based methods [15-17]. In the deep learning methods, the geometric and atomic information of sample structures were used to extract features, and the actual protein-ligand binding sites were used as training labels. The mentioned deep learning models were brought out in the past two years. These models were basically derived from UNet, whereas PUResNet adopted ResNet to integrate the encoder side of UNet [15-19]. Residual neural network is known as ResNet, which displayed higher accuracy due to the increased depth can better extract target features. ResNet was developed to cope with the problem of vanishing gradient by introducing skip connection and summing subsequent layers [19]. The introducing of ResNet to PUResNet has increased the depth of the model, which contributed to a better performance than the only UNet based model Kalasanty [16]. However, ResNet showed inherent limitations due to the skip connection can affect the learning potential of the network. This may explain the minor improved accuracy of PUResNet compared with Kalasanty. Deeppocket organized a two-step protocol. Firstly, Fpocket was used to score the potential binding sites in the protein-ligand complex, then the top ranked binding sites were passed to Deeppocket for searching the potential ligand binding sites [7, 15].

Inspired by the previous studies [15-17], in this work, we mainly focused on rebuilding the CNN model using a more accurate model, and enlarging the training set to further optimize the model performance. Previous reports adopted different versions of scPDB dataset for training the deep learning model [15-17, 20]. Classification analysis of the scPDB datasets based on the Uniprot ID showed that the scPDB dataset contain less than 5540 non-reductant protein structures in the previous studies [15, 17]. Due to the size of training set can affect the accuracy of CNN models [21], we considered the size of scPDB is not satisfied for training. We enlarged the training set by gaining novel samples or rotating the protein and cavity structures by 15 degrees in the original scPBD_5020 dataset (derived scPDB dataset developed in the study of PUResNet) [16, 22]. These datasets were independently used to train DUnet.

DUnet was developed based on the framework of DenseNet and UNet. DenseNet emerges more recent than ResNet, and displayed better performance [23]. DenseNet adopts dense concatenation to the subsequent layers so that the features of each layers are maintained. Recently, the utility of DenseNet in solving protein-based tasks has highlighted its performance in extracting features of 3D targets [24]. In this study, DenseNet was used to solve the 3D image segmentation problem. DUnet was trained and compared with the current state-of-art method PUResNet [16]. The performance of DUnet was checked used the validation sets and our built industrial enzyme dataset. The predicted binding sites of microbial transglutaminase were experimentally validated their contribution to enzyme activity by Ala scan. On the basis of model accuracy, DUnet was used to predict the ligand binding sites of Swiss-Prot annotated proteins [25].

## Materials and methods

### Computational procedure

#### Data preparation

Protein structures were converted into a 3D grid at a size of 70 Å^3^. The grid was represented as voxels at a size of 2 Å^3^ to ensure each voxel can only contain one atom (Fig. S1). Protein structures were described according to the atomic physical characters, and converted to tensors for training the model (Fig. S1). Each heavy atom was described by 18 atomic features, such as atom types, hybridization, in bound with other heavy atoms or heteroatoms, and partial charge [26]. The output array is in a shape of 36×36×36×1, where atomic features was replaced by a value of 1 and 0 to indicate the ligand binding or non-binding sites, respectively. For training labels, the ligand binding sites were extracted as array with an initial shape of 36×36×36×18, and the shape is changed to 36×36×36×1 by replacing the atomic feature by a value of 1 or 0 to represent if the given voxel contains binding sites.

Previous study highlighted the importance of using enlarged dataset to optimize the model performance [21, 27]. In this work, we prepared three training sets with different sample sizes: (1) scPDB_5020 dataset (totally 5020 protein-ligand samples) [16]; (2) scPDB_5020 and SC6K_C datasets (totally 6833 protein-ligand samples) [15, 16]; (3) scPDB_5020 and scPDB5020_r datasets (totally 10040 protein-ligand samples), and the resulted trained model were named DUnet-1, DUnet-2, and DUnet-3. The protein data in scPDB_5020 dataset was cleaned the water molecule, metals, and ligands before uses. The SC6K_C dataset was prepared by removing 677 structures that is unreadable or displayed a distance between ligand and protein surface is more than 6 Å, and the protein data was cleaned as mentioned above. Inspired by the data argumentation method, we prepared scPDB5020_r dataset by rotating the protein and cavity structures in the scPDB_5020 by 15 degrees [28] (Fig. S2). Considering the size of protein grid is not able to cover the full protein, the input features can be changed by rotating the protein structure. Here, rotating the structure by 15 degrees is an empirical value.

The validation sets COACH420 and BU48 contained 298 and 62 protein structures, respectively [16]. We built a small dataset contains 11 commonly used industrial enzymes and their substrates or inhibitors for validation purpose. The Swiss-Prot annotated protein structures were downloaded from AlphaFold Protein Structure Database, and we removed structure file less than 100 kb since these structures are single alpha helix with loops which are not formal structures.

#### Model architecture

DUnet framework was derived from DenseNet and UNet for investigating comprehensive protein-ligand binding sites problems. In terms of Unet architecture, DUnet contains an encoder and decoder side, where encoder side in charge of extracting image features through convolution, and decoder side is used for recovering the image size. The Dense block was mainly integrated in the encoder side. The encoder and decoder side both contain five major blocks as indicated in Fig. 1. These main blocks were mainly consist of sub-blocks including Dense, Transition, and Upsampling block (Fig. 1). Dense, Transition, and Upsampling block consist of 39, 6, and 5 layers separately (Fig. 1), and the detailed information is provided in Additional file 1. Concatenation, which also known as “skip connection” was used to fast-forward high-resolution feature maps from the encoder to the decoder side, and benefit the model convergence (Fig. 1). Sigmoid activation function was used in the decoder part to finally translate the point value to 0 and 1 to match the training labels. Dice loss was used to solve the segmentation problem in this study. DUnet was trained using three independent dataset for 100 epochs with the optimal batch size and learning rate. DUnet was built by PyTorch with 17 727 969 trainable parameters. We adopted Adam optimizer, and optimized the batch size and learning rate. The model was trained using NVIDIA 3080TI GPU hardware for 100 epochs. The reference model PUResNet was downloaded accordingly [16], and we adopted 2 methods for evaluating and comparing the model accuracy.

**Figure 1.**
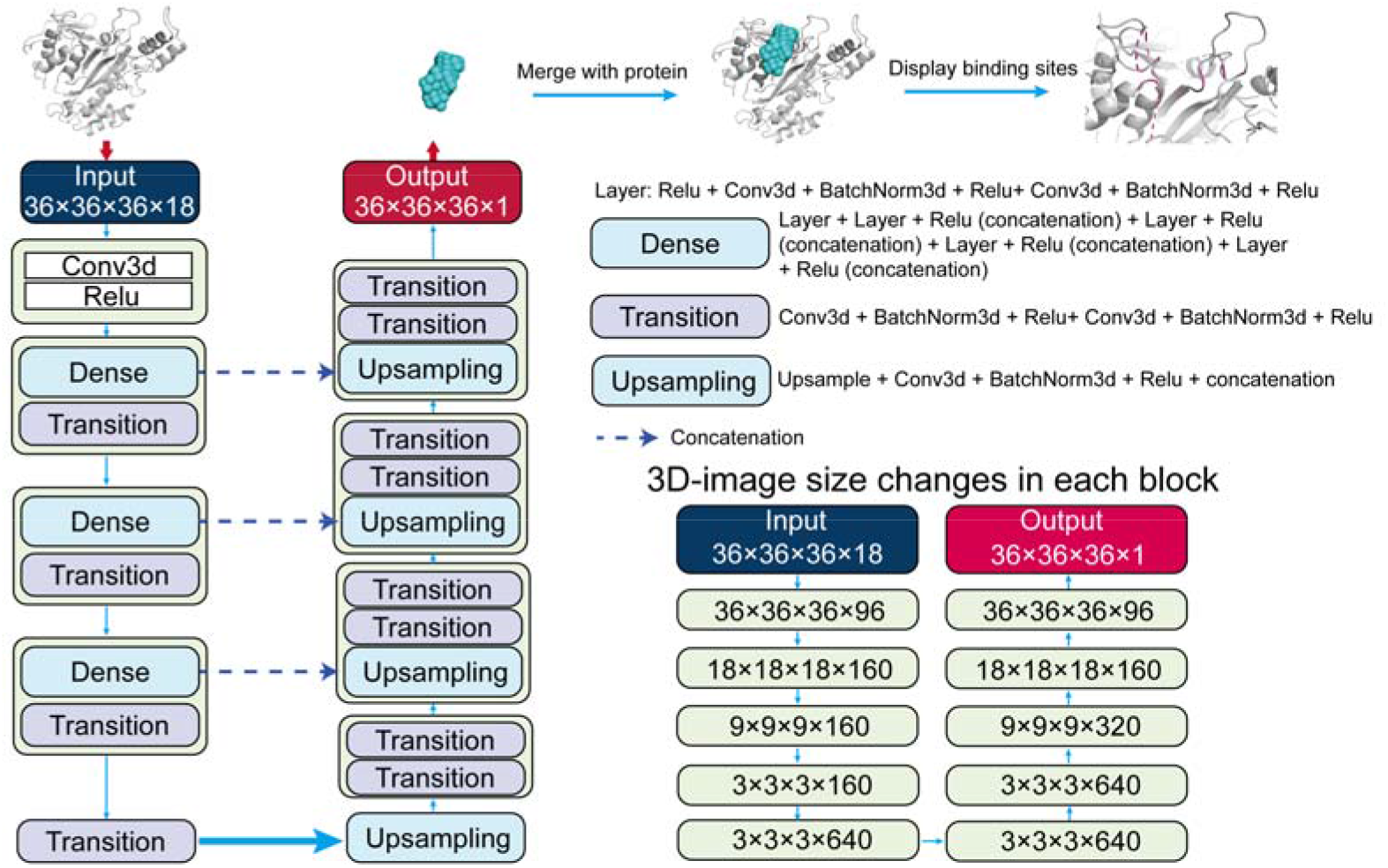
Architecture of DUnet.

#### Model evaluation

Distance center-center (DCC) was used to measure the barycenter distance between the predicted binding sites and actual ligand; discretized volume overlap (DVO) was used to judge the voxels overlap between the predicted binding sites and actual ligand (Fig. S3). Here, we developed a strict rule for DVO calculations rather than using flattening method as the dice loss function [16, 17]. The coordinates of the predicted binding sites and actual ligand were rounded, and evaluated the 3D-points match (Fig. S3).

### Experimental procedure

#### Strains and plasmids

The plasmid pET-22b (+) and *Escherichia coli* (*E. coli*) BL21 (DE3) were used for expressing *Streptomyces mobaraenesis* transglutaminase (smTG) and its variants.

#### Plasmid construction

The gene encoding for smTG carried its native pro-peptide (GenBank accession number: EU301664) was synthesized and cloned into pET-22b via *Nde*I and *Blp*I sites, yielding the plasmids pET-22b/*smTG* (GENEWIZ, Suzhou, China). The plasmids encoding the smTG with single residue mutations were constructed by polymerase chain reaction (PCR) using pET-22b/*smTG* as template, and the primers were listed in Table S1. The PCR products were purified and circularized using Blunting Kination Ligation Kit (TaKaRa, Dalian, China)

#### Protein expression and purification

The plasmid encoding for smTG and its variants were transformed into *E. coli*, and the obtained single colony was inoculated into Luria-Bertani (LB) medium supplemented with 50 µg/mL ampicillin. The seed culture was cultivated at 37□ for 10 h, and was transferred into Terrifc-broth (TB) supplemented with 50 µg/mL ampicillin and cultivated until the cell density (OD_600_) reached to 1.0. Isopropylthio-β-galactoside (IPTG) was then supplemented to 0.1 mM for the induction of recombinant expressed proteins. The strain was cultivated under 20□ for 30 h.

Cells were obtained from the fermentation culture by centrifugation, and resuspended in Tris-HCl (50 mM, pH 8.0) for ultra-sonification. The pro-peptide of smTG and its variants was removed using dispase (Solarbio, Beijing, China) at a final concentration of 2 mg/mL and incubated at 37□ for 30 min. The obtained solutions were subjected to affinity chromatography using His-Trap column (GE Healthcare, New York, USA) and size-exclusion chromatography (SEC) using Superdex 75 column (GE Healthcare, New York, USA). Finally, smTG and its variants were in Tris-HCl (50 mM, pH 8.0) for enzyme analysis.

Protein concentration was determined using Bradford Protein Assay Kit (Beyotime, Shanghai, China). Sodium dodecyl sulfate-polyacrylamide gel electrophoresis (SDS-PAGE) was conducted using 12% Tris-glycine gel (ThermoFisher, Shanghai, China).

#### Determination of transglutaminase activity assay

To measure the specific activity of smTG, 60 μL of pre-warmed (at 37□ for 5 min) sample protein was added to 150 μL substrate solution (200 mM Tris-HCl, 100 mM hydroxylamine, 10 mM GSH, 30 mM CBZ-Gln-Gly, pH 6.0), and the reaction was terminated by 60 μL of termination solution (termination solution achieved by mixing same volume of 3 M HCl, 12% trichloroacetic acid and 5% FeCl_3_·H_2_O) after 10 min incubation under 37□ [29]. One unit of smTG activity was defined as 1 μmol L-glutamic acid γ-monohydroxamate produced per min.

## Results and discussion

### Model performance on validation sets

#### K-fold cross-validation

For initial estimating the accuracy of DUnet, scPDB_5020 dataset was randomly split into 5-fold for cross-validation [17]. We use 4 folds for training the model, and 1 fold for validation. With an optimal batch size of 5 and learning rate of 0.001, the average loss for the 5 folds was 27% (Fig. S4). The result DCC ≤ 4 Å was defined as success according to previous studies [16, 17]. In this study, if multiple binding sites were predicted we took only the first one into account. The average success rate of the 5-fold was 79% (Table S2). The 2^nd^ fold displayed the highest success rate which reached 84% (Table S2).

#### DCC analysis

Firstly, we compared the DCC ≤ 4 Å results for each model. On the COACH420 validation set, the DCC ≤ 4 Å for PUResNet, DUnet-1, DUnet-2, and DUnet-3 were 49.6%, 43.3%, 43.6% and 45% (Table 1), respectively. Here, if the model predicts multiple pockets, only the first predicted pocket was used. This explained the DCC ≤ 4 Å result from PUResNet is lower than the reported 51% [16]. On the BU48 validation set, the four models displayed 51.6%, 50%, 51.6% and 66.1% (Table 1), respectively.

**Table 1.** Performance of PUResNet and DUnet on validation set

| | | $DCC \leq 4\text{\AA}$ | DVO<br>($DCC \leq 4\text{\AA}$) | DVO<br>(DCC<br>top-ranked<br>150) | DVO<br>(DCC<br>top-ranked<br>31) | Failed |
| --- | --- | --- | --- | --- | --- | --- |
| COACH420 | PUNet | 49.6% | 0.020 | 0.020 | - | 18 |
|  | DUNet-1 | 43.3% | 0.030 | 0.028 | - | 0 |
|  | DUNet-2 | 43.6% | 0.030 | 0.030 | - | 0 |
|  | DUNet-3 | 45% | 0.030 | 0.028 | - | 0 |
| BU48 | PUNet | 51.6% | 0.021 | - | 0.022 | 0 |
|  | DUNet-1 | 50% | 0.026 | - | 0.029 | 1 |
|  | DUNet-2 | 51.6% | 0.028 | - | 0.031 | 0 |
|  | DUNet-3 | 66.1% | 0.027 | - | 0.031 | 0 |

\*DUNet-1: DUNet trained with scPDB\_5020 dataset
\*DUNet-2: DUNet trained with combined scPDB\_5020 and SC6K\_C dataset
\*DUNet-3: DUNet trained with combined scPDB\_5020 and scPDB5020\_r dataset
\*Failed: Failed to predict any binding regions

Obviously, DUnet trained with larger dataset contributes to higher values of DCC ≤ 4 Å, especially for DUnet-3 which was trained using the largest dataset generated by data argumentation. However, DUnet-2 trained with scPDB_5020 and SC6K_C datasets only gave minor increases.

DUnet-3 displayed dramatic improvement in DCC (Table 1). It increased the value of DCC ≤ 4 Å by 1.7% compared with DUnet simply trained using scPDB_5020 dataset, and it increased the value on BU48 validation set was 30% higher more than that of predicted by PUResNet and DUnet trained using the other two datasets (Fig. 2). Due to DUnet-3 used the enlarged dataset is consist of scPDB_5020 and scPDB5020_r which achieved by rotating all the proteins and ligands in the scPDB_5020 dataset by 15°, we checked data argumentation was not simply doubled the sample size based on scPDB_5020. The top-ranked 100 DCC results on COACH420 validation set achieved by DUnet-1 and DUnet-3 showed only 64 Protein Data Bank (PDB) IDs overlaps (Fig. 3). This is lower than that of predicted by PUResNet and DUnet-1, which has a value of 74 (Fig. 3). Moreover, the DCC for PDB ID: 2cx8 achieved by DUnet-1 and DUnet-3 were 0.22 and 27 Å, respectively. Therefore, the influence of using data argumentation approach is significant.

**Figure 2.**
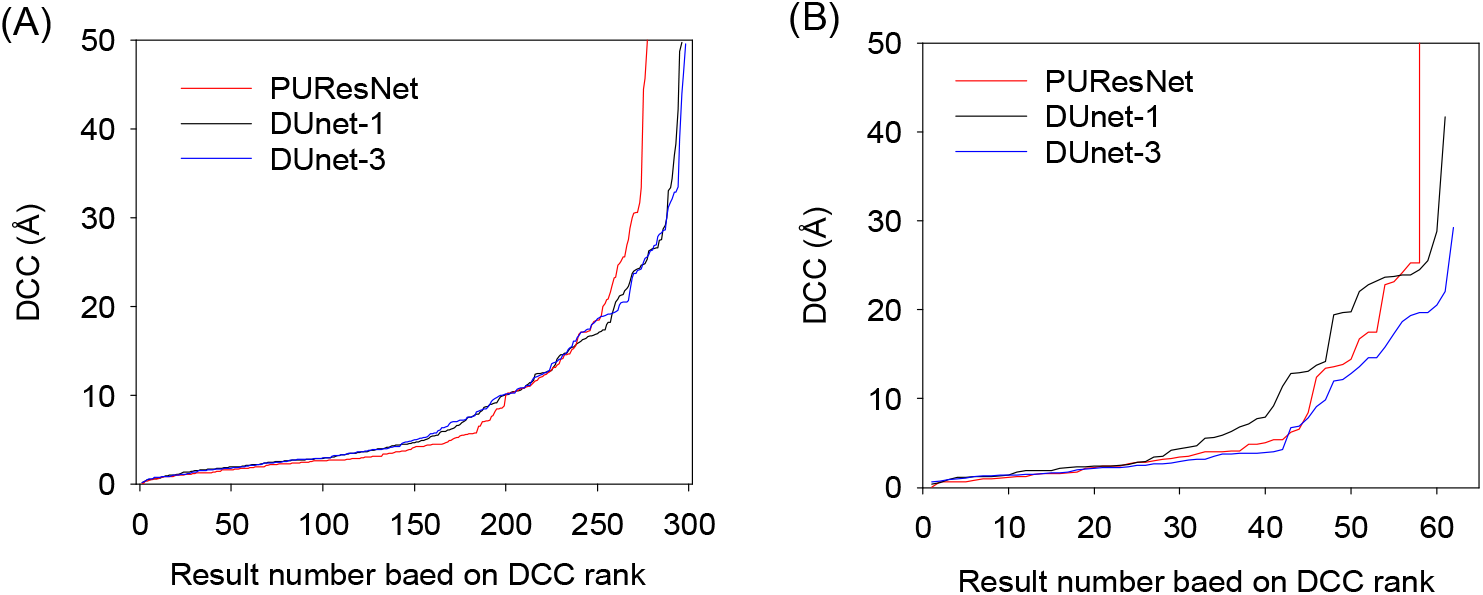
DCC result alignment on validation set. (A) DCC result of PUResNet and DUnet on COACH420 validation set; (B) DCC result of PUResNet and DUnet on BU48 validation set.

**Figure 3.**
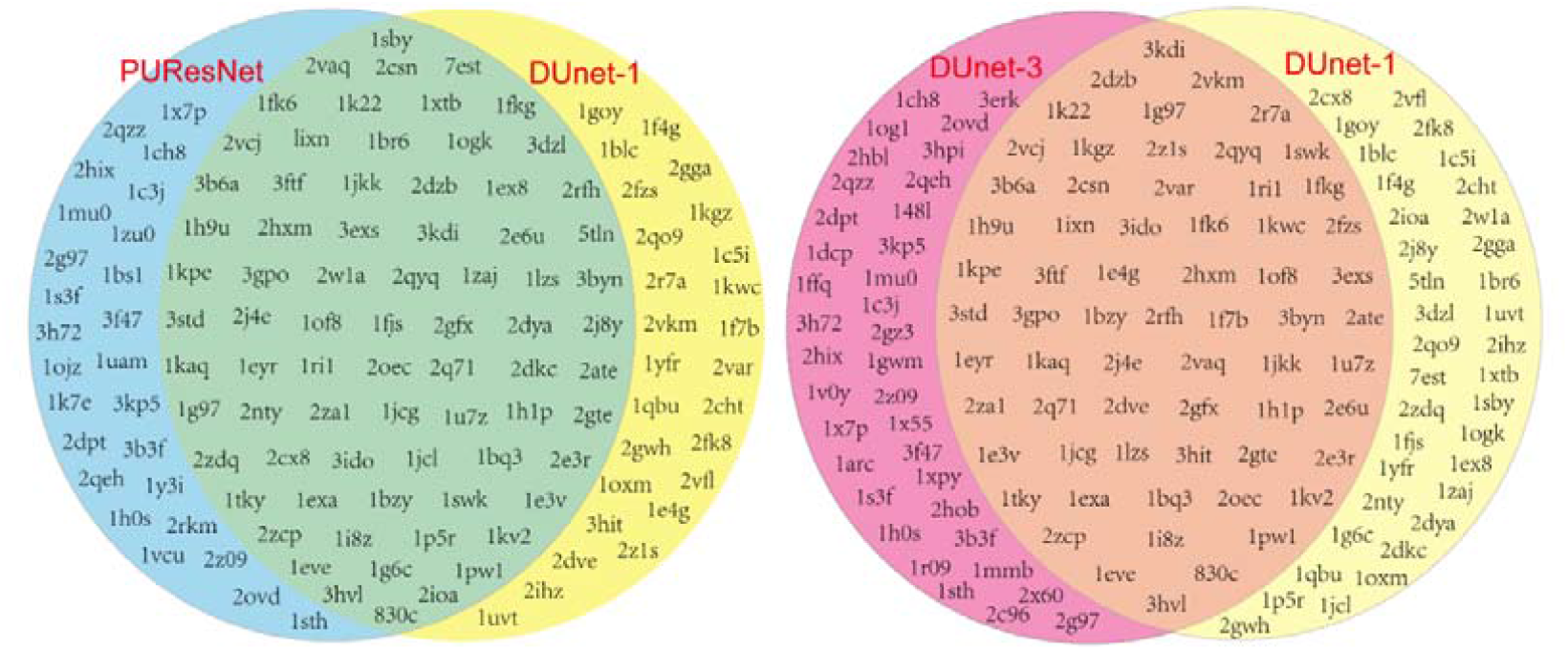
The common predicted results in DCC top-ranked 100 predicted by PUResNet and DUnet on the COACH420 validation set.

However, it seems the performance using DCC evaluations of DUnet-1 and DUnet-2 were worse than that of PUResNet. To clarify this phenomenon with technical analysis, we visually inspected the DCC top-ranked 100 on COACH420 validation set predicted by PUResNet and DUnet-1 which trained on the same dataset (Additional File 2-4). We noticed that the predicted ligand binding sites using PUResNet were much larger than that predicted by DUnet-1, which contained more non-ligand areas (Additional File 2-4). As mentioned above, the training labels in scPDB_5020 dataset are ligands binding sites without non-binding area. But, if predicted ligand area is far larger than the real ligand area, it will certainly more easy to cover the actual binding areas, and achieve lower DCC as it provide closed barycenter center. The examples for can be found in Additional file 2-4. The predicted large ligand binding sites can also lead to lower DVO, which was described in the DVO analysis section.

Visual inspection of the top-ranked 100 DCC result achieved by PUResNet and DUnet-1 highlighted that DCC ≤ 4 Å as the only evaluation method for judging success rates is not sufficient. We agree with Marta *et al*., that using DVO to represent the intersection of the predicted and the true ligand position is a more stringent method [17]. Obtaining high DVO requires not only the predicted location is accurate, but also the predicted ligand binding sites are not excessively larger than the actual ligand. Therefore, higher DVO is usually difficult to obtain [16, 17].

#### DVO analysis

Because predictions by PUResNet and DUnet gave different DCC results, comparison of DVO only based on DCC ≤ 4 Å results will lead to unequal sample sizes. In addition to comparing the DVO for DCC ≤ 4 Å, we also compared the DVO of the DCC top-ranked 150 results predicted by PUResNet and DUnet. As shown in Table 1, all versions of DUnet displayed higher DVO values than PUResNet, even DUnet-1 has increased the value by more than 24% (Table 1). Both DUnet-2 and DUnet-3 showed higher DVO than DUnet-1, which indicated that enlarging training set could also contribute to accurate predicting of the ligand-binding horizon (Table 1). However, comparing DVO values achieved by DUnet-2 and DUnet-3 showed that the data argument strategy not further improve the accuracy, while resulting in minor decreases (Table 1).

Due to the DVO method used in this study aims to align the point matches of predicted binding sites and actual ligand coordinates, and the ligand is not with hydrogen bonds, the DVO value calculated for all the models are far smaller than the reported values in other studies [15-17]. Through visual inspecting the results, we noticed that even the DVO values were relative low, but the predicted ligand already covered all protein-binding sites and the redundant area is within a reasonable range of intervals (Additional File 2-4). In other words, the achieved value by DUnet may already be a high value that a model can reach.

At last, DUnet dramatically reduced the failed numbers during predictions compared to PUResNet. The failed number for PUResNet and DUnet-1 on validation set COACH420 were 18 and 0, and for BU48 the number was and 0 and 1 (Table 1).

### Predicting enzyme-substrate binding sites for industrial enzymes

Enzymes are important bio-catalysts widely used for food processing and chemical synthesis to feed industrial needs [30]. Searching for substrate binding area is the key to select “hot spots” for engineering enzyme activity. In this study, we aim to prove that DUnet can be used for locating the enzyme-substrates binding sites. We collected 11 enzyme-substrate structures from PDB, including cutinase, lipase, alpha-amylase, beta-amylase, trypsin, cutinase, alkaline proteinase, xylanase, cellulase, beta-1,4-endoglucanase, and transglutaminase (Table 2), and architected dataset EN11.

**Table 2.** Data information in dataset EN11

| PDB_ID | Protein name | Organism |
| --- | --- | --- |
| 1oa7 | Cellulase | *Melanocarpus albomyces* |
| 1pig | Alpha-Amylase | *Sus scrofa* |
| 1r50 | Lipase | *Bacillus subtilis* |
| 1scn | Alkaline proteinase | *Bacillus licheniformis* |
| 1q6c | Beta-Amylase | *Glycine max* |
| 1v2w | Trypsin | *Bos taurus* |
| 2bnj | Xylanase | *Thermoascus aurantiacus* |
| 3esb | Cutinase | *Fusarium vanettenii* |
| 4pse | Cutinase | *Trichoderma reesei QM6a* |
| 5i79 | Beta-1,4-endoglucanase | *Aspergillus niger* |
| 6gmg | Transglutaminase | *Streptomyces mobaraenesis* |

As shown in Fig. 4A, the number of DCC ≤ 4 Å achieved by PUResNet and DUnet were 5 and 6, and the average DVO of DCC ≤ 4 Å achieved by the two models was the same (0.033). PUResNet failed to predict the binding sites for alkaline proteinase (PDB ID: 1scn). Through visualizing the predicted binding sites, we found the two models showed similar results despite of their DCC differences (Fig. 4B). We were unsure if these deep learning-based models have better performances compared to traditional physics-based method. So, we used FTSite (up-to-date version) which scores the cavity mainly based on the binding affinity between target protein and virtual ligands as a reference model [8, 31]. We aligned the predicted results achieved by FTSite with PUResNet and DUnet on EN11. In FTSite, the predicted binding area with pink color has the highest priority (Fig. 4B). Through visual inspection, we observed that FTSite gave wrong prediction for 1oa7, for PUResNet were 1scn and 1v2w (rank 1 was wrong), and for DUnet-3 was 1r50. Visual inspect suggests that DUnet-3 also provided better coverage on the ligand binding sites than the other two models.

**Figure 4.**
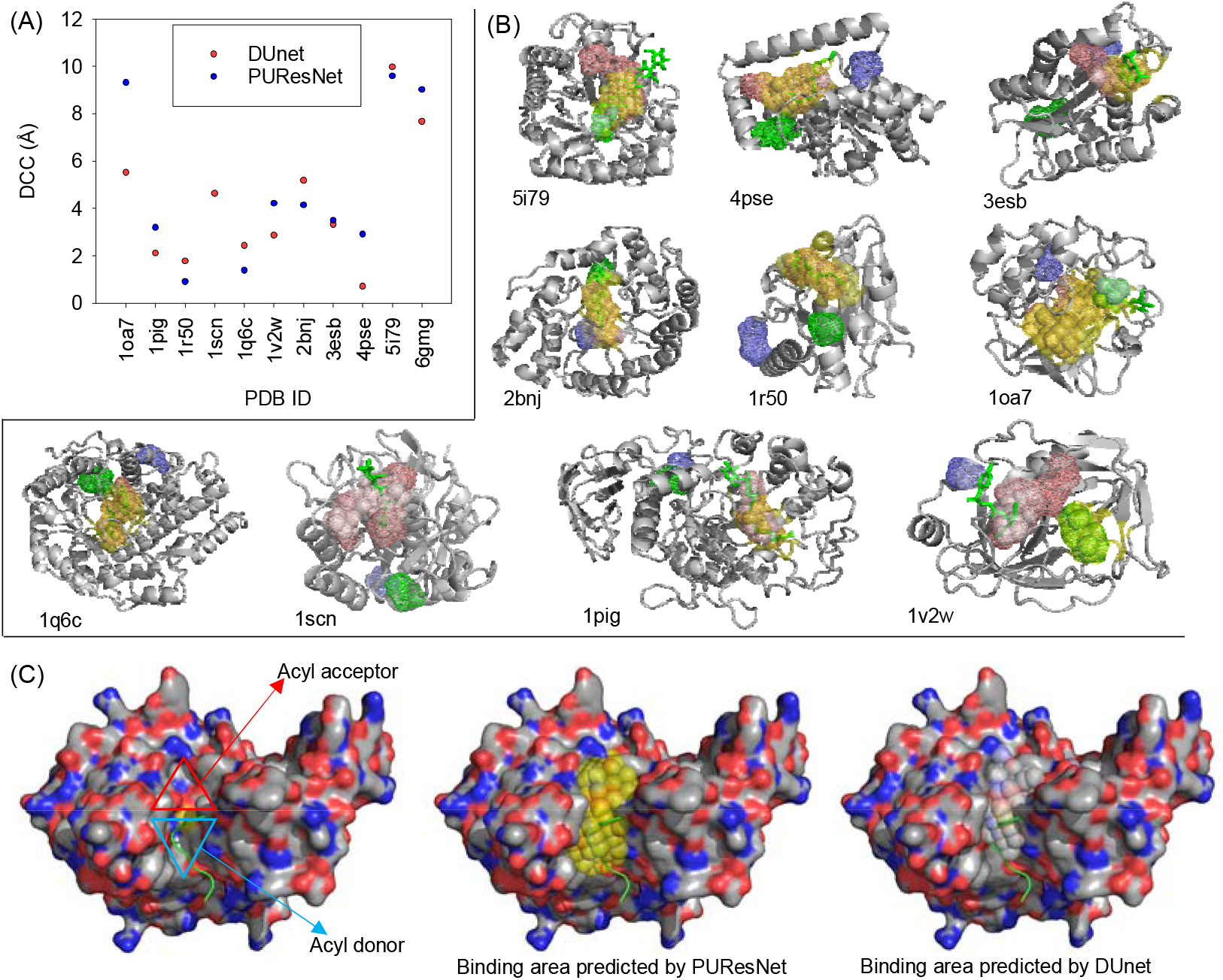
PUResNet and DUnet on the EN11 validation set. (A) DCC result of PUResNet and DUnet-3 on EN11 validation set; (B) Visualization of predicted binding sites by FTSite, PUResNet, and DUnet-3. For FTSite prediction, the mesh representation in pink was the most recommended prediction; the sphere representation in yellow was predicted region by PUResNet, and in white was predicted by DUnet-3; (C) The predicted binding sites of transglutaminase (PDB ID: 6gmg) by PUResNet and DUnet-3, the substrate position of acyl donor and acceptor was presented by triangle, and the cartoon representation in green color is the acyl donor. The sphere representation in yellow was predicted region by PUResNet, and in white was predicted region by DUnet-3.

### Experimentally validating predicting binding sites on transglutaminase

Our team has previously worked on the modification of transglutaminase (smTG) derived from *Streptomyces mobaraenesis* (PDB ID: 6gmg), this enzyme contains an acyl donor and acceptor binding area within its catalytic pocket (Fig. 4C) [32]. In the given structure (PDB ID: 6gmg), only acyl donor was co-crystallized while the acyl acceptor was absent. Previous study has addressed the importance of acyl acceptor binding sites to the enzyme activity [33]. To the best of the authors’ knowledge, the acyl acceptor binding sites were not fully characterized. DUnet-3 predicts 28 key residues for both acyl donor and acceptor binding, while 19 residues suggested to influence the acyl donor binding were previous characterized by Ala scan [34]. In this study, we further characterized the 9 residues that mainly responsible for acyl acceptor binding by Ala scan (Fig. 5A). smTG and its variants were purified and visualized on SDS-PAGE, which showed these enzymes are in high purity (Fig. 5B). The obtained activity for smTG was 24 U/mg (Fig. 5C). Our result showed that mutants V290A, Y291A, and E292A displayed no activity, and mutants T273A, G275A, and W298A reduced their activity by more than 40% (Fig. 5C). Only mutants H289A, N297A, and N301A remained more than 75% activity (Fig. 5C). Considering 66.7% of the selected sites displayed obvious reduced activity, we concluded the predicted acyl acceptor binding sites was correct.

**Figure 5.**
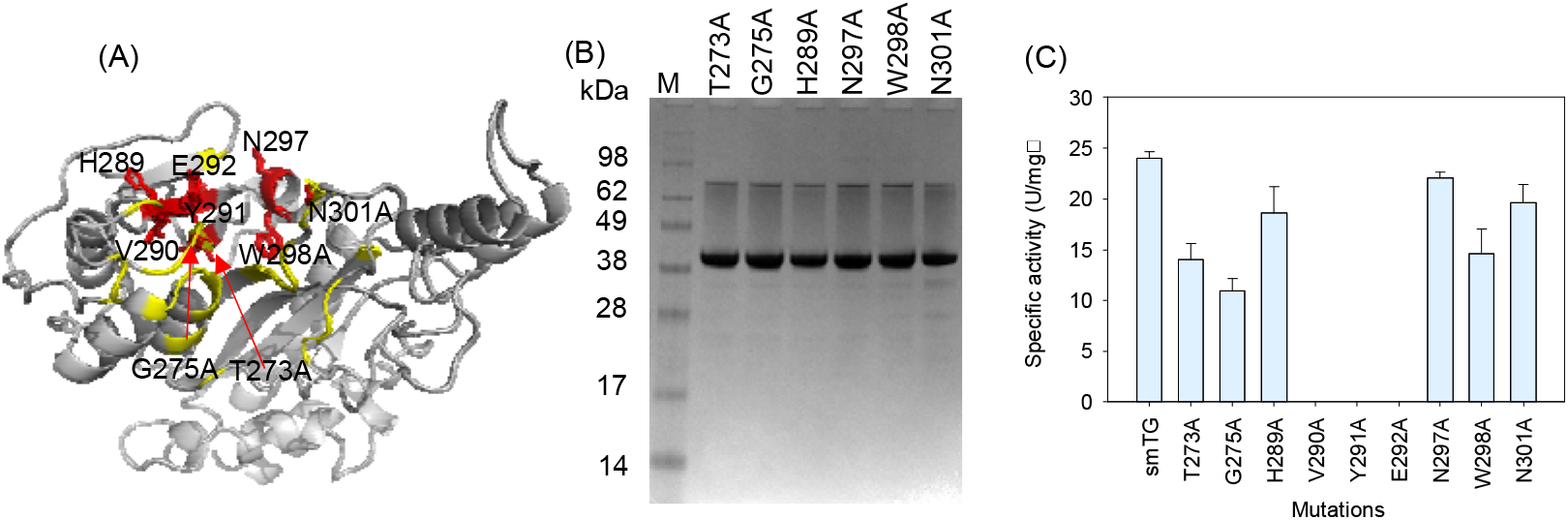
Characterization the predicted binding sites for transglutaminase. (A) Stereo-view of experimentally tested sites in this study, the tested sites were in red, and previous tested sites were represented in yellow [34]; (B) Visualization of purified smTG and its variants on SDS-PAGE; (C) Specific activity of smTG and its variants.

### Potential uses of DUnet

Until the manuscript was finished, the deposited structures in PDB were approximately 190 thousand, which is still limited structural information considering natural existing proteins. Previously, AlphaFold-2 has predicted structures for Swiss-Prot annotated proteins, which can be public accessed through AlphaFold Protein Structure Database and Uniprot [25]. Currently, the Swiss-Prot dataset contained 542,380 protein data entries. The proteins were majorly originated from bacteria, which have more than 330 thousand entries, whereas eukaryotic and archaeal have over 190 thousand and 19 thousand entries. The predicted structures have provided convenience for protein engineering, and widely used for designing novel drugs or metabolic pathway-related importance enzymes. In order to satisfy the need for localizing the key enzyme-substrate binding sites, we made predictions of Swiss-Prot annotated proteins using DUnet-3. For convenient uses, all the predicted results in were deposited in GitHub for public access. We provided the predicted ligand binding sites and the exact binding residues within proteins. The protein-ligand binding sites were conducted by searching the neighbor residues within a distance of 4 Å of virtual ligands. These results provide support for the protein/enzyme engineering and benefit the activity design protocol.

The use of DenseNet has already contributed to the protein design and drug developing tasks [24, 35]. In this work, we architected DUnet that derived from UNet and DenseNet. We showed that DUnet is capable for protein and ligand binding sites segmentation. This framework can be used to train other tasks, such as evaluating mutations related to protein stability, recognition or clustering of small molecules, and *de novo* design structures.

## Conclusion

In the present study, we introduced a 3D-CNN based model DUnet for predicting the protein-ligand binding sites. The model was trained using scPDB dataset, and enhanced its accuracy by enlarging the training set. We adopted data argumentation strategy for enlarging the training set, which showed great contribution to the prediction of the barycenter of predicted and actual ligands. The model was validated and aligned with the current state-of-art methods, which displayed better performance. DUnet also provided accurate predictions for enzyme-substrate binding sites. We experimentally validated its predictions on the binding sites of microbial transglutaminase, and showed the significance of these sites. These results suggested that DUnet can be a priory solution for locating protein-ligand binding sites. Based on the performance, DUnet was used to predict the binding area of Swiss-Prot annotated proteins to enable convenient uses. DUnet can serve the protein activity engineering protocol, and provide a general framework for other tasks.

## Data availability

The supplementary material is uploaded in Supplemented.docx

Additional file 1: The layer, shape, and parameter information of DUnet.

Additional file 2: Commonly existed 64 PDB IDs in DCC top-ranked 100 predicted by PUResNet and DUnet-1 on COACH420 validation set. The substrate presented in stick representation, the sphere representation in yellow color was predicted ligand region by PUResNet, and in white color was predicted ligand region by DUnet-1.

Additional file 3: Except the commonly existed PDB IDs in the DCC top-ranked 100 results predicted by DUnet-1.

Additional file 4: Except the commonly existed PDB IDs in the DCC top-ranked 100 results predicted by PUResNet.

## Code availability

All the codes and datasets used this work are publicly available at: https://github.com/wangxinglong1990/DUnet.

## Acknowledgements

This work was supported by the National Key Research and Development Program of China (No. 2019YFA0706900) and the National Natural Science Foundation of China (No. 32071474 and 31771913).

## Author information

## Contributions

BZ and XW: coding. PY, YT, RM, JD, and XW: Additional material processing. JZ, SR, JC, SL, and XW: supervising the project. BZ and XW: manuscript writing. XW: results analysis. All authors read and approved the final manuscript.

## Reference

1. Walker SP, Yallapragada VVB, Tangney M (2021) Arming Yourself for The In Silico Protein Design Revolution. Trends Biotechnol 39(7):651–664. https://doi.org/10.1016/j.tibtech.2020.10.003

2. Jumper J, Evans R, Pritzel A, Green T, Figurnov M, Ronneberger O, Tunyasuvunakool K, Bates R, Žídek A, Potapenko A, Bridgland A, Meyer C, Kohl SAA, Ballard AJ, Cowie A, Romera-Paredes B, Nikolov S, Jain R, Adler J, Back T, Petersen S, Reiman D, Clancy E, Zielinski M, Steinegger M, Pacholska M, Berghammer T, Bodenstein S, Silver D, Vinyals O, Senior AW, Kavukcuoglu K, Kohli P, Hassabis D (2021) Highly accurate protein structure prediction with AlphaFold. Nature 596(7873):583–589. https://doi:10.1038/s41586-021-03819-2

3. Baek M, DiMaio F, Anishchenko I, Dauparas J, Ovchinnikov S, Lee GR, Wang J, Cong Q, Kinch LN, Schaeffer RD, Millán C, Park H, Adams C, Glassman CR, DeGiovanni A, Pereira JH, Rodrigues AV, Dijk AAv, Ebrecht AC, Opperman DJ, Sagmeister T, Buhlheller C, Pavkov-Keller T, Rathinaswamy MK, Dalwadi U, Yip CK, Burke JE, Garcia KC, Grishin NV, Adams PD, Read RJ, Baker D (2021) Accurate prediction of protein structures and interactions using a three-track neural network. Science 373(6557):871–876. https://doi:10.1126/science.abj8754

4. Park H, Bradley P, Greisen P, Jr., Liu Y, Mulligan VK, Kim DE, Baker D, DiMaio F (2016) Simultaneous Optimization of Biomolecular Energy Functions on Features from Small Molecules and Macromolecules. J Chem Theory Comput 12(12):6201–6212. https://doi:10.1021/acs.jctc.6b00819

5. Buß O, Rudat J, Ochsenreither K (2018) FoldX as Protein Engineering Tool: Better Than Random Based Approaches? Computational and Structural Biotechnology Journal 16(25-33. https://doi.org/10.1016/j.csbj.2018.01.002

6. Zhou H, Cao H, Skolnick J (2018) FINDSITEcomb2.0: A New Approach for Virtual Ligand Screening of Proteins and Virtual Target Screening of Biomolecules. J Chem Inf Model 58(11):2343–2354.https://doi:10.1021/acs.jcim.8b00309

7. Schmidtke P, Le Guilloux V, Maupetit J, Tufféry P (2010) fpocket: online tools for protein ensemble pocket detection and tracking. Nucleic Acids Res 38(Web Server issue):W582–W589. https://doi:10.1093/nar/gkq383

8. Ngan C-H, Hall DR, Zerbe B, Grove LE, Kozakov D, Vajda S (2012) FTSite: high accuracy detection of ligand binding sites on unbound protein structures. Bioinformatics 28(2):286–287. https://doi:10.1093/bioinformatics/btr651

9. Huang B, Schroeder M (2006) LIGSITEcsc: predicting ligand binding sites using the Connolly surface and degree of conservation. BMC Structural Biology 6(1):19. https://doi.org/10.1186/1472-6807-6-19

10. Xie Z-R, Hwang MJ (2012) Ligand-binding site prediction using ligand-interacting and binding site-enriched protein triangles. Bioinformatics 28(12):1579–1585. https://doi.org/10.1093/bioinformatics/bts182

11. He K, Gkioxari G, Dollár P, Girshick R (2017) Mask R-CNN. In 2017 IEEE International Conference on Computer Vision (ICCV); 22-29 Oct. 2017. 2980–2988.

12. Dietler N, Minder M, Gligorovski V, Economou AM, Joly DAHL, Sadeghi A, Chan CHM, Koziński M, Weigert M, Bitbol A-F, Rahi SJ (2020) A convolutional neural network segments yeast microscopy images with high accuracy. Nat Commun 11(1):5723. https://doi:10.1038/s41467-020-19557-4

13. Liu Z, Su W, Ao J, Wang M, Jiang Q, He J, Gao H, Lei S, Nie J, Yan X, Guo X, Zhou P, Hu H, Ji M (2022) Instant diagnosis of gastroscopic biopsy via deep-learned single-shot femtosecond stimulated Raman histology. Nat Commun 13(1):4050. https://doi.org/10.1038/s41467-022-31339-8

14. Hänsch A, Chlebus G, Meine H, Thielke F, Kock F, Paulus T, Abolmaali N, Schenk A (2022) Improving automatic liver tumor segmentation in late-phase MRI using multi-model training and 3D convolutional neural networks. Sci Rep 12(1):12262. https://doi:10.1038/s41598-022-16388-9

15. Aggarwal R, Gupta A, Chelur V, Jawahar CV, Priyakumar UD (2021) DeepPocket: Ligand Binding Site Detection and Segmentation using 3D Convolutional Neural Networks. J Chem Inf Model. https://doi:10.1021/acs.jcim.1c00799

16. Kandel J, Tayara H, Chong KT (2021) PUResNet: prediction of protein-ligand binding sites using deep residual neural network. J Cheminformatics 13(1):65. https://doi:10.1186/s13321-021-00547-7

17. Stepniewska-Dziubinska MM, Zielenkiewicz P, Siedlecki P (2020) Improving detection of protein-ligand binding sites with 3D segmentation. Sci Rep 10(1):5035. https://doi:10.1038/s41598-020-61860-z

18. Ronneberger O, Fischer P, Brox T (2015) U-Net: Convolutional Networks for Biomedical Image Segmentation. In Medical Image Computing and Computer-Assisted Intervention – MICCAI 2015; Cham. Edited by Navab N, Hornegger J, Wells WM, Frangi AF. Springer International Publishing: 234–241.

19. He K, Zhang X, Ren S, Sun J (2015) Deep Residual Learning for Image Recognition. arXiv. https://doi.org/10.48550/arXiv.1512.03385

20. Meslamani J, Rognan D, Kellenberger E (2011) sc-PDB: a database for identifying variations and multiplicity of ‘druggable’ binding sites in proteins. Bioinformatics 27(9):1324–1326. https://doi.org/10.1093/bioinformatics/btr120

21. Ng W, Minasny B, Mendes WDS, Demattê JAM (2020) The influence of training sample size on the accuracy of deep learning models for the prediction of soil properties with near-infrared spectroscopy data. Soil 6(2):565–578. https://doi:10.5194/soil-6-565-2020

22. Monshi MMA, Poon J, Chung V, Monshi FM (2021) CovidXrayNet: Optimizing data augmentation and CNN hyperparameters for improved COVID-19 detection from CXR. Comput Biol Med 133(104375-104375. https://doi:10.1016/j.compbiomed.2021.104375

23. Huang G, Liu Z, Maaten LVD, Weinberger KQ (2017) Densely Connected Convolutional Networks. In 2017 IEEE Conference on Computer Vision and Pattern Recognition (CVPR); 21-26 July 2017. 2261–2269.

24. Qi Y, Zhang JZH (2020) DenseCPD: Improving the Accuracy of Neural-Network-Based Computational Protein Sequence Design with DenseNet. J Chem Inf Model 60(3):1245–1252. https://doi:10.1021/acs.jcim.0c00043

25. Consortium TU (2018) UniProt: a worldwide hub of protein knowledge. Nucleic Acids Res 47(D1):D506–D515. https://doi:10.1093/nar/gky1049

26. Stepniewska-Dziubinska MM, Zielenkiewicz P, Siedlecki P (2018) Development and evaluation of a deep learning model for protein-ligand binding affinity prediction. Bioinformatics 34(21):3666–3674. https://doi:10.1093/bioinformatics/bty374

27. Jiang J, Liu H, Zhao C, He C, Ma J, Cheng T, Zhu Y, Cao W, Yao X (2022) Evaluation of Diverse Convolutional Neural Networks and Training Strategies for Wheat Leaf Disease Identification with Field-Acquired Photographs. Remote Sens 14(14):3446. https://doi:10.3390/rs14143446

28. Van Der Spoel D, Lindahl E, Hess B, Groenhof G, Mark AE, Berendsen HJC (2005) GROMACS: Fast, flexible, and free. Journal of Computational Chemistry 26(16):1701–1718. https://doi.org/10.1002/jcc.20291

29. Wang X, Du J, Zhao B, Wang H, Rao S, Du G, Zhou J, Chen J, Liu S (2021) Significantly Improving the Thermostability and Catalytic Efficiency of Streptomyces mobaraenesis Transglutaminase through Combined Rational Design. Journal of Agricultural and Food Chemistry 69(50):15268–15278. https://doi.org/10.1021/acs.jafc.1c05256

30. Bornscheuer UT, Huisman GW, Kazlauskas RJ, Lutz S, Moore JC, Robins K (2012) Engineering the third wave of biocatalysis. Nature 485(7397):185–194. https://doi:10.1038/nature11117

31. Kozakov D, Grove LE, Hall DR, Bohnuud T, Mottarella SE, Luo L, Xia B, Beglov D, Vajda S (2015) The FTMap family of web servers for determining and characterizing ligand-binding hot spots of proteins. Nat Protoc 10(5):733–755. https://doi:10.1038/nprot.2015.043

32. Juettner NE, Schmelz S, Kraemer A, Knapp S, Becker B, Kolmar H, Scrima A, Fuchsbauer H-L (2018) Structure of a glutamine donor mimicking inhibitory peptide shaped by the catalytic cleft of microbial transglutaminase. FEBS J 285(24):4684–4694. https://doi.org/10.1111/febs.14678

33. Oteng-Pabi SK, Keillor JW (2013) Continuous enzyme-coupled assay for microbial transglutaminase activity. Analytical Biochemistry 441(2):169–173. https://doi.org/10.1016/j.ab.2013.07.014

34. Tagami U, Shimba N, Nakamura M, Yokoyama K-i, Suzuki E-i, Hirokawa T (2009) Substrate specificity of microbial transglutaminase as revealed by three-dimensional docking simulation and mutagenesis. Protein Engineering, Design and Selection 22(12):747–752. https://doi.org/10.1093/protein/gzp061

35. Li Y, Hu J, Wang Y, Zhou J, Zhang L, Liu Z (2020) DeepScaffold: A Comprehensive Tool for Scaffold-Based De Novo Drug Discovery Using Deep Learning. J Chem Inf Model 60(1):77–91. https://doi.org/10.1021/acs.jcim.9b00727

