## Supplementary figures and images for "DUnet: A deep learning guided protein-ligand binding pocket prediction"

### Additional file 2

#### Slide 1
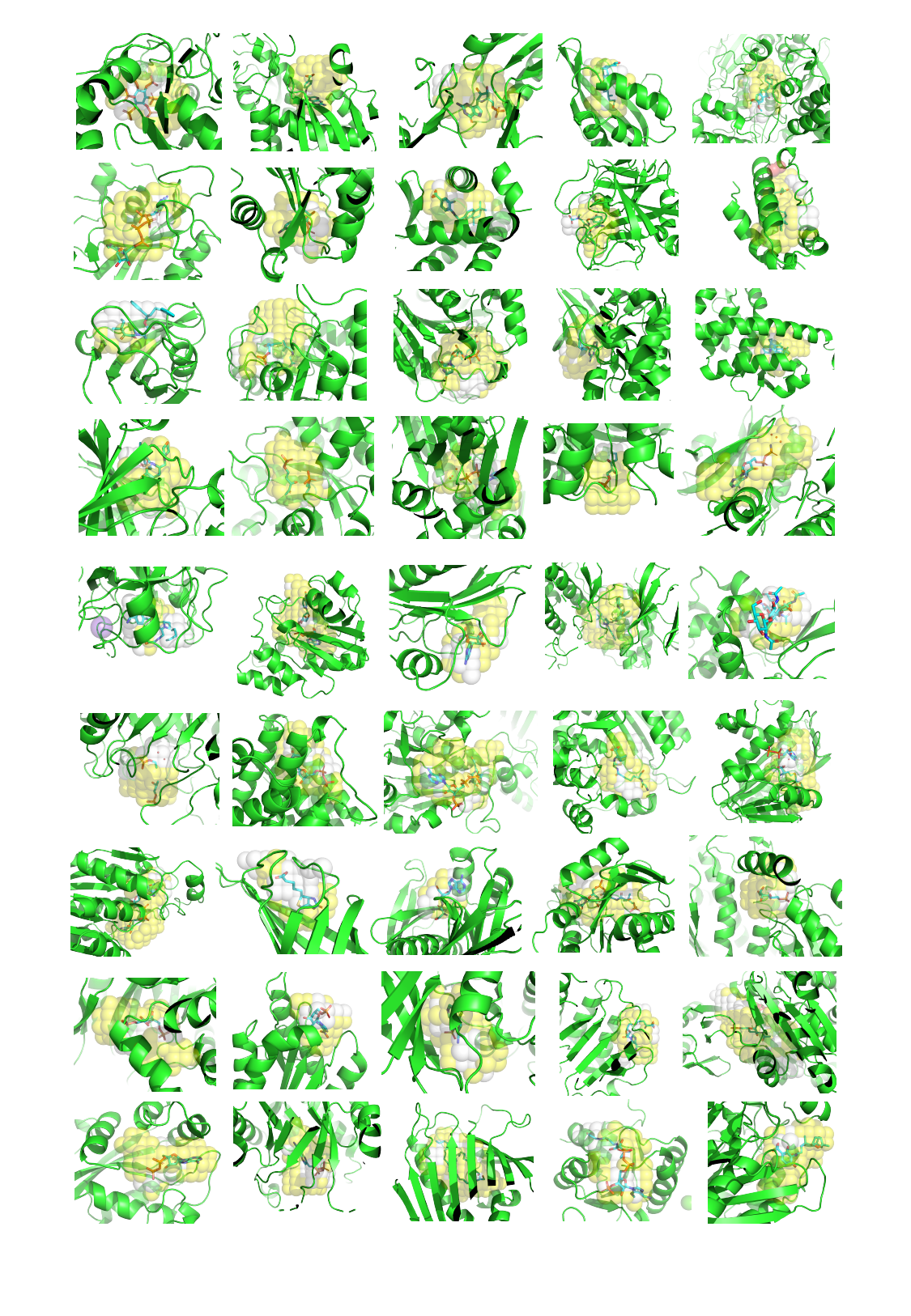

#### Slide 2
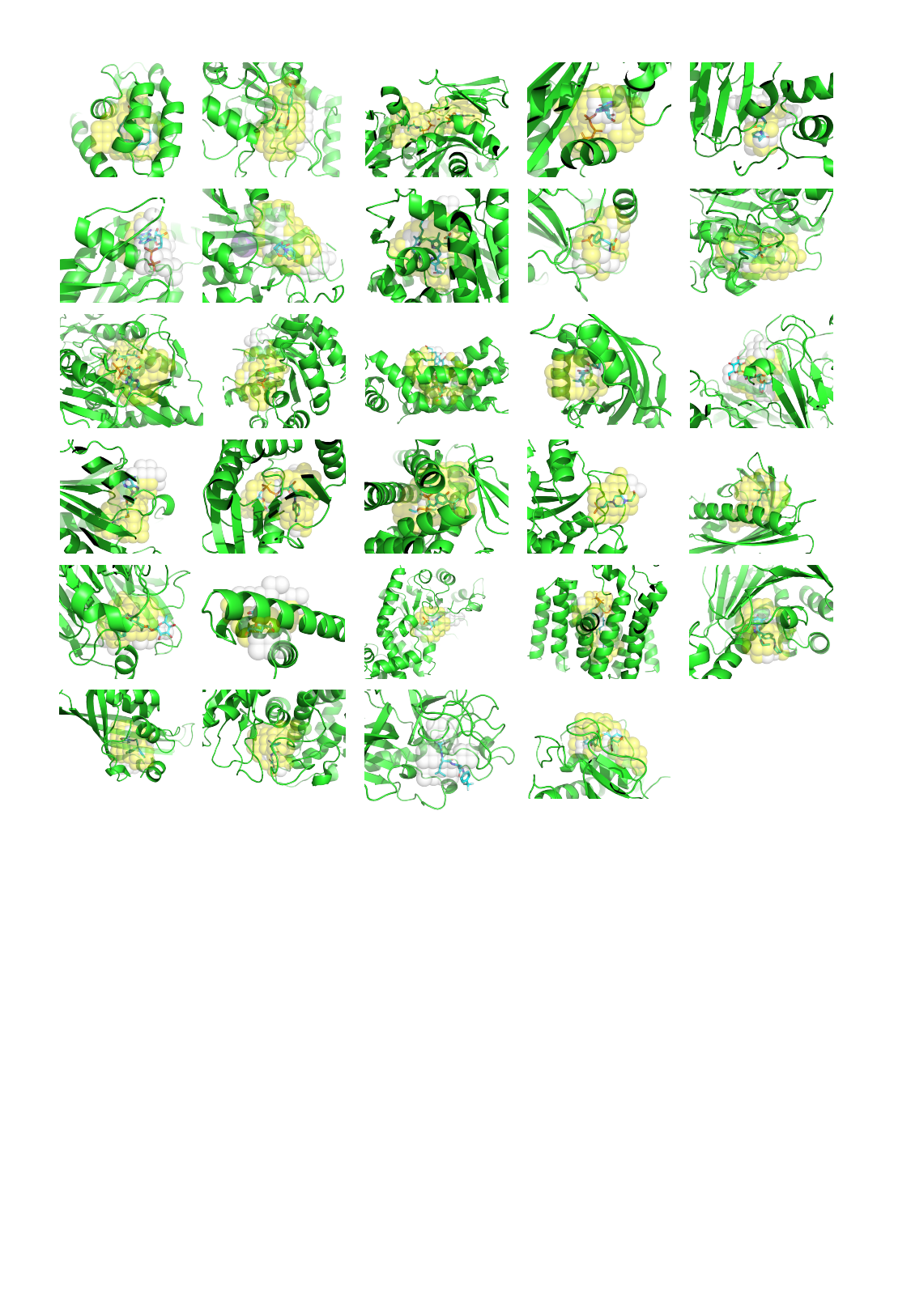

### Additional file 3

#### Slide 1
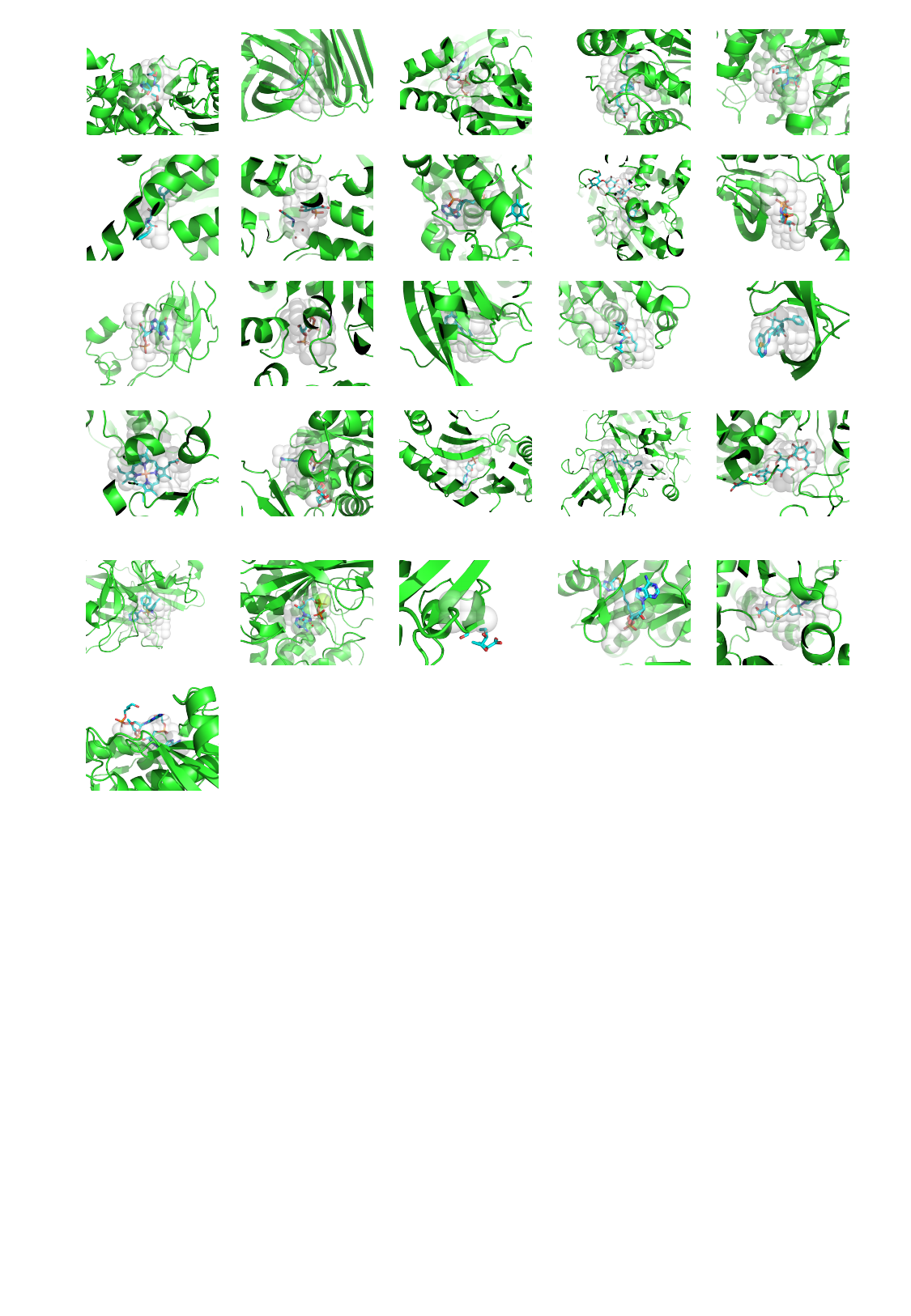

### Additional file 4

#### Slide 1
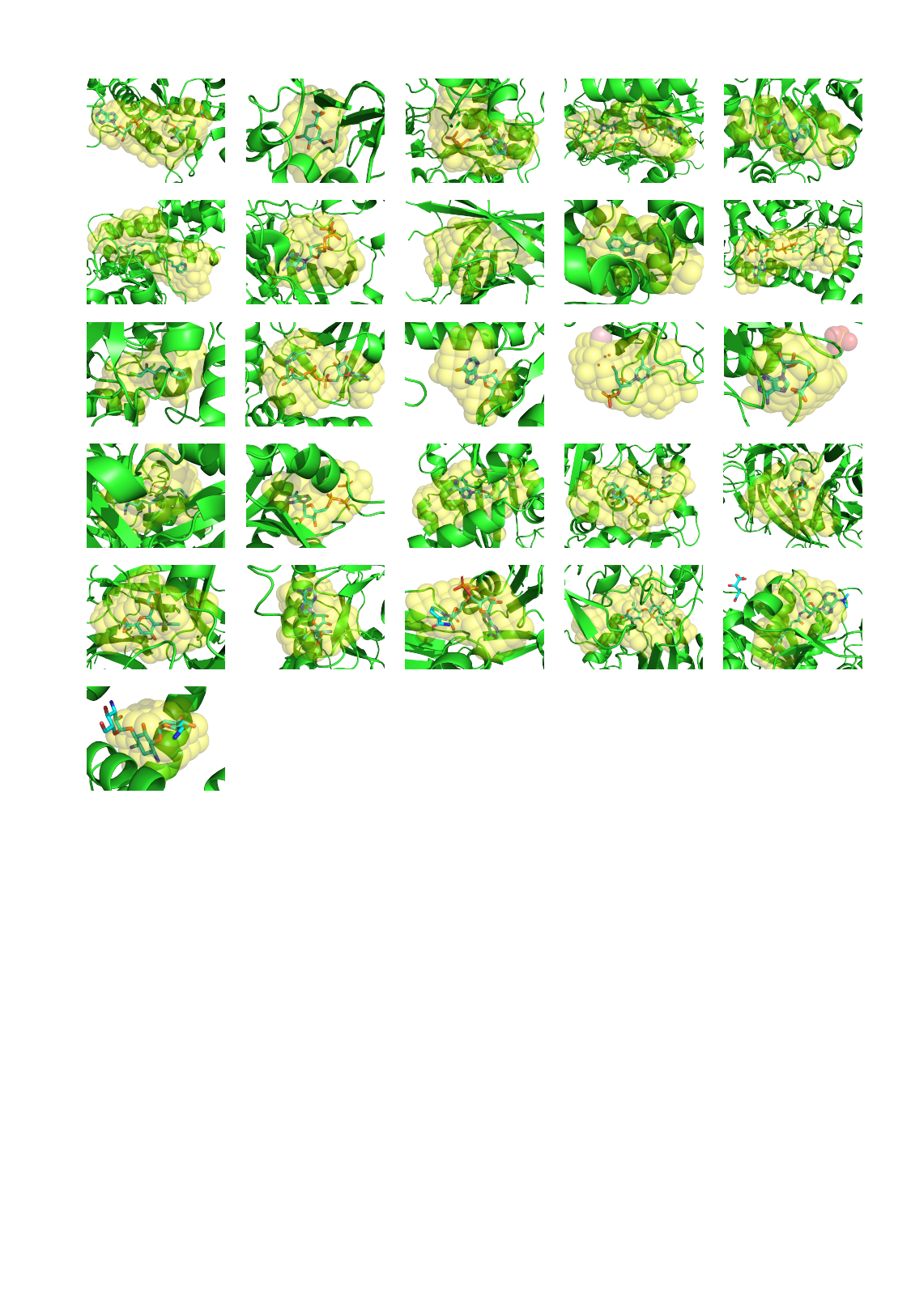
