## Additional file 1 for "DUnet: A deep learning guided protein-ligand binding pocket prediction"

| Layer | Shape | parameters |
| --- | --- | --- |
| Conv3d-1 | [-1,96,36,36,36] | 592,800 |
| BatchNorm3d-2 | [-1,96,36,36,36] | 192 |
| ReLU-3 | [-1,96,36,36,36] | 0 |
| Conv3d-4 | [-1,96,36,36,36] | 9,312 |
| BatchNorm3d-5 | [-1,96,36,36,36] | 192 |
| ReLU-6 | [-1,96,36,36,36] | 0 |
| Conv3d-7 | [-1,96,36,36,36] | 9,312 |
| BatchNorm3d-8 | [-1,96,36,36,36] | 192 |
| ReLU-9 | [-1,96,36,36,36] | 0 |
| TransitionBlock-10 | [-1,96,36,36,36] | 0 |
| Conv3d-11 | [-1,96,36,36,36] | 9,312 |
| BatchNorm3d-12 | [-1,96,36,36,36] | 192 |
| ReLU-13 | [-1,96,36,36,36] | 0 |
| Conv3d-14 | [-1,96,18,18,18] | 9,312 |
| BatchNorm3d-15 | [-1,96,18,18,18] | 192 |
| ReLU-16 | [-1,96,18,18,18] | 0 |
| TransitionBlock-17 | [-1,96,18,18,18] | 0 |
| BatchNorm3d-18 | [-1,96,18,18,18] | 192 |
| Conv3d-19 | [-1,128,18,18,18] | 12,416 |
| ReLU-20 | [-1,128,18,18,18] | 0 |
| BatchNorm3d-21 | [-1,128,18,18,18] | 256 |
| Conv3d-22 | [-1,32,18,18,18] | 110,624 |
| ReLU-23 | [-1,32,18,18,18] | 0 |
| BatchNorm3d-24 | [-1,32,18,18,18] | 64 |
| Conv3d-25 | [-1,128,18,18,18] | 4,224 |
| ReLU-26 | [-1,128,18,18,18] | 0 |
| BatchNorm3d-27 | [-1,128,18,18,18] | 256 |
| Conv3d-28 | [-1,32,18,18,18] | 110,624 |
| ReLU-29 | [-1,32,18,18,18] | 0 |
| ReLU-30 | [-1,64,18,18,18] | 0 |
| BatchNorm3d-31 | [-1,64,18,18,18] | 128 |
| Conv3d-32 | [-1,128,18,18,18] | 8,320 |
| ReLU-33 | [-1,128,18,18,18] | 0 |
| BatchNorm3d-34 | [-1,128,18,18,18] | 256 |
| Conv3d-35 | [-1,32,18,18,18] | 110,624 |
| ReLU-36 | [-1,32,18,18,18] | 0 |
| ReLU-37 | [-1,96,18,18,18] | 0 |
| BatchNorm3d-38 | [-1,96,18,18,18] | 192 |
| Conv3d-39 | [-1,128,18,18,18] | 12,416 |
| ReLU-40 | [-1,128,18,18,18] | 0 |
| BatchNorm3d-41 | [-1,128,18,18,18] | 256 |
| Conv3d-42 | [-1,32,18,18,18] | 110,624 |
| ReLU-43 | [-1,32,18,18,18] | 0 |
| ReLU-44 | [-1,128,18,18,18] | 0 |
| BatchNorm3d-45 | [-1,128,18,18,18] | 256 |
| Conv3d-46 | [-1,128,18,18,18] | 16,512 |
| ReLU-47 | [-1,128,18,18,18] | 0 |
| BatchNorm3d-48 | [-1,128,18,18,18] | 256 |
| Conv3d-49 | [-1,32,18,18,18] | 110,624 |
| ReLU-50 | [-1,32,18,18,18] | 0 |
| ReLU-51 | [-1,160,18,18,18] | 0 |
| DenseBlock-52 | [-1,160,18,18,18] | 0 |
| Conv3d-53 | [-1,160,18,18,18] | 25,760 |
| BatchNorm3d-54 | [-1,160,18,18,18] | 320 |
| ReLU-55 | [-1,160,18,18,18] | 0 |
| Conv3d-56 | [-1,160,18,18,18] | 25,760 |
| BatchNorm3d-57 | [-1,160,18,18,18] | 320 |
| ReLU-58 | [-1,160,18,18,18] | 0 |
| TransitionBlock-59 | [-1,160,18,18,18] | 0 |
| Conv3d-60 | [-1,160,18,18,18] | 25,760 |
| BatchNorm3d-61 | [-1,160,18,18,18] | 320 |
| ReLU-62 | [-1,160,18,18,18] | 0 |
| Conv3d-63 | [-1,160,9,9,9] | 25,760 |
| BatchNorm3d-64 | [-1,160,9,9,9] | 320 |
| ReLU-65 | [-1,160,9,9,9] | 0 |
| TransitionBlock-66 | [-1,160,9,9,9] | 0 |
| BatchNorm3d-67 | [-1,160,9,9,9] | 320 |
| Conv3d-68 | [-1,128,9,9,9] | 20,608 |
| ReLU-69 | [-1,128,9,9,9] | 0 |
| BatchNorm3d-70 | [-1,128,9,9,9] | 256 |
| Conv3d-71 | [-1,32,9,9,9] | 110,624 |
| ReLU-72 | [-1,32,9,9,9] | 0 |
| BatchNorm3d-73 | [-1,32,9,9,9] | 64 |
| Conv3d-74 | [-1,128,9,9,9] | 4,224 |
| ReLU-75 | [-1,128,9,9,9] | 0 |
| BatchNorm3d-76 | [-1,128,9,9,9] | 256 |
| Conv3d-77 | [-1,32,9,9,9] | 110,624 |
| ReLU-78 | [-1,32,9,9,9] | 0 |
| ReLU-79 | [-1,64,9,9,9] | 0 |
| BatchNorm3d-80 | [-1,64,9,9,9] | 128 |
| Conv3d-81 | [-1,128,9,9,9] | 8,320 |
| ReLU-82 | [-1,128,9,9,9] | 0 |
| BatchNorm3d-83 | [-1,128,9,9,9] | 256 |
| Conv3d-84 | [-1,32,9,9,9] | 110,624 |
| ReLU-85 | [-1,32,9,9,9] | 0 |
| ReLU-86 | [-1,96,9,9,9] | 0 |
| BatchNorm3d-87 | [-1,96,9,9,9] | 192 |
| Conv3d-88 | [-1,128,9,9,9] | 12,416 |
| ReLU-89 | [-1,128,9,9,9] | 0 |
| BatchNorm3d-90 | [-1,128,9,9,9] | 256 |
| Conv3d-91 | [-1,32,9,9,9] | 110,624 |
| ReLU-92 | [-1,32,9,9,9] | 0 |
| ReLU-93 | [-1,128,9,9,9] | 0 |
| BatchNorm3d-94 | [-1,128,9,9,9] | 256 |
| Conv3d-95 | [-1,128,9,9,9] | 16,512 |
| ReLU-96 | [-1,128,9,9,9] | 0 |
| BatchNorm3d-97 | [-1,128,9,9,9] | 256 |
| Conv3d-98 | [-1,32,9,9,9] | 110,624 |
| ReLU-99 | [-1,32,9,9,9] | 0 |
| ReLU-100 | [-1,160,9,9,9] | 0 |
| DenseBlock-101 | [-1,160,9,9,9] | 0 |
| Conv3d-102 | [-1,160,9,9,9] | 25,760 |
| BatchNorm3d-103 | [-1,160,9,9,9] | 320 |
| ReLU-104 | [-1,160,9,9,9] | 0 |
| Conv3d-105 | [-1,160,9,9,9] | 25,760 |
| BatchNorm3d-106 | [-1,160,9,9,9] | 320 |
| ReLU-107 | [-1,160,9,9,9] | 0 |
| TransitionBlock-108 | [-1,160,9,9,9] | 0 |
| Conv3d-109 | [-1,160,9,9,9] | 25,760 |
| BatchNorm3d-110 | [-1,160,9,9,9] | 320 |
| ReLU-111 | [-1,160,9,9,9] | 0 |
| Conv3d-112 | [-1,160,3,3,3] | 25,760 |
| BatchNorm3d-113 | [-1,160,3,3,3] | 320 |
| ReLU-114 | [-1,160,3,3,3] | 0 |
| TransitionBlock-115 | [-1,160,3,3,3] | 0 |
| BatchNorm3d-116 | [-1,160,3,3,3] | 320 |
| Conv3d-117 | [-1,128,3,3,3] | 20,608 |
| ReLU-118 | [-1,128,3,3,3] | 0 |
| BatchNorm3d-119 | [-1,128,3,3,3] | 256 |
| Conv3d-120 | [-1,32,3,3,3] | 110,624 |
| ReLU-121 | [-1,32,3,3,3] | 0 |
| BatchNorm3d-122 | [-1,32,3,3,3] | 64 |
| Conv3d-123 | [-1,128,3,3,3] | 4,224 |
| ReLU-124 | [-1,128,3,3,3] | 0 |
| BatchNorm3d-125 | [-1,128,3,3,3] | 256 |
| Conv3d-126 | [-1,32,3,3,3] | 110,624 |
| ReLU-127 | [-1,32,3,3,3] | 0 |
| ReLU-128 | [-1,64,3,3,3] | 0 |
| BatchNorm3d-129 | [-1,64,3,3,3] | 128 |
| Conv3d-130 | [-1,128,3,3,3] | 8,320 |
| ReLU-131 | [-1,128,3,3,3] | 0 |
| BatchNorm3d-132 | [-1,128,3,3,3] | 256 |
| Conv3d-133 | [-1,32,3,3,3] | 110,624 |
| ReLU-134 | [-1,32,3,3,3] | 0 |
| ReLU-135 | [-1,96,3,3,3] | 0 |
| BatchNorm3d-136 | [-1,96,3,3,3] | 192 |
| Conv3d-137 | [-1,128,3,3,3] | 12,416 |
| ReLU-138 | [-1,128,3,3,3] | 0 |
| BatchNorm3d-139 | [-1,128,3,3,3] | 256 |
| Conv3d-140 | [-1,32,3,3,3] | 110,624 |
| ReLU-141 | [-1,32,3,3,3] | 0 |
| ReLU-142 | [-1,128,3,3,3] | 0 |
| BatchNorm3d-143 | [-1,128,3,3,3] | 256 |
| Conv3d-144 | [-1,128,3,3,3] | 16,512 |
| ReLU-145 | [-1,128,3,3,3] | 0 |
| BatchNorm3d-146 | [-1,128,3,3,3] | 256 |
| Conv3d-147 | [-1,32,3,3,3] | 110,624 |
| ReLU-148 | [-1,32,3,3,3] | 0 |
| ReLU-149 | [-1,160,3,3,3] | 0 |
| DenseBlock-150 | [-1,160,3,3,3] | 0 |
| Conv3d-151 | [-1,320,3,3,3] | 51,520 |
| BatchNorm3d-152 | [-1,320,3,3,3] | 640 |
| ReLU-153 | [-1,320,3,3,3] | 0 |
| Conv3d-154 | [-1,640,3,3,3] | 205,440 |
| BatchNorm3d-155 | [-1,640,3,3,3] | 1,280 |
| ReLU-156 | [-1,640,3,3,3] | 0 |
| Upsample-157 | [-1,640,3,3,3] | 0 |
| Upsample-158 | [-1,640,3,3,3] | 0 |
| Conv3d-159 | [-1,640,3,3,3] | ######## |
| BatchNorm3d-160 | [-1,640,3,3,3] | 1,280 |
| ReLU-161 | [-1,640,3,3,3] | 0 |
| UpsamplingBlock-162 | [-1,640,3,3,3] | 0 |
| Conv3d-163 | [-1,640,3,3,3] | 410,240 |
| BatchNorm3d-164 | [-1,640,3,3,3] | 1,280 |
| ReLU-165 | [-1,640,3,3,3] | 0 |
| Conv3d-166 | [-1,640,3,3,3] | 410,240 |
| BatchNorm3d-167 | [-1,640,3,3,3] | 1,280 |
| ReLU-168 | [-1,640,3,3,3] | 0 |
| TransitionBlock-169 | [-1,640,3,3,3] | 0 |
| Conv3d-170 | [-1,640,3,3,3] | 410,240 |
| BatchNorm3d-171 | [-1,640,3,3,3] | 1,280 |
| ReLU-172 | [-1,640,3,3,3] | 0 |
| Conv3d-173 | [-1,640,3,3,3] | 410,240 |
| BatchNorm3d-174 | [-1,640,3,3,3] | 1,280 |
| ReLU-175 | [-1,640,3,3,3] | 0 |
| TransitionBlock-176 | [-1,640,3,3,3] | 0 |
| Upsample-177 | [-1,640,9,9,9] | 0 |
| Upsample-178 | [-1,160,9,9,9] | 0 |
| Conv3d-179 | [-1,320,9,9,9] | ######## |
| BatchNorm3d-180 | [-1,320,9,9,9] | 640 |
| ReLU-181 | [-1,320,9,9,9] | 0 |
| UpsamplingBlock-182 | [-1,320,9,9,9] | 0 |
| Conv3d-183 | [-1,320,9,9,9] | 102,720 |
| BatchNorm3d-184 | [-1,320,9,9,9] | 640 |
| ReLU-185 | [-1,320,9,9,9] | 0 |
| Conv3d-186 | [-1,320,9,9,9] | 102,720 |
| BatchNorm3d-187 | [-1,320,9,9,9] | 640 |
| ReLU-188 | [-1,320,9,9,9] | 0 |
| TransitionBlock-189 | [-1,320,9,9,9] | 0 |
| Conv3d-190 | [-1,320,9,9,9] | 102,720 |
| BatchNorm3d-191 | [-1,320,9,9,9] | 640 |
| ReLU-192 | [-1,320,9,9,9] | 0 |
| Conv3d-193 | [-1,320,9,9,9] | 102,720 |
| BatchNorm3d-194 | [-1,320,9,9,9] | 640 |
| ReLU-195 | [-1,320,9,9,9] | 0 |
| TransitionBlock-196 | [-1,320,9,9,9] | 0 |
| Upsample-197 | [-1,320,18,18,18] | 0 |
| Upsample-198 | [-1,160,18,18,18] | 0 |
| Conv3d-199 | [-1,160,18,18,18] | ######## |
| BatchNorm3d-200 | [-1,160,18,18,18] | 320 |
| ReLU-201 | [-1,160,18,18,18] | 0 |
| UpsamplingBlock-202 | [-1,160,18,18,18] | 0 |
| Conv3d-203 | [-1,160,18,18,18] | 25,760 |
| BatchNorm3d-204 | [-1,160,18,18,18] | 320 |
| ReLU-205 | [-1,160,18,18,18] | 0 |
| Conv3d-206 | [-1,160,18,18,18] | 25,760 |
| BatchNorm3d-207 | [-1,160,18,18,18] | 320 |
| ReLU-208 | [-1,160,18,18,18] | 0 |
| TransitionBlock-209 | [-1,160,18,18,18] | 0 |
| Conv3d-210 | [-1,160,18,18,18] | 25,760 |
| BatchNorm3d-211 | [-1,160,18,18,18] | 320 |
| ReLU-212 | [-1,160,18,18,18] | 0 |
| Conv3d-213 | [-1,160,18,18,18] | 25,760 |
| BatchNorm3d-214 | [-1,160,18,18,18] | 320 |
| ReLU-215 | [-1,160,18,18,18] | 0 |
| TransitionBlock-216 | [-1,160,18,18,18] | 0 |
| Upsample-217 | [-1,160,36,36,36] | 0 |
| Upsample-218 | [-1,160,36,36,36] | 0 |
| Conv3d-219 | [-1,96,36,36,36] | 829,536 |
| BatchNorm3d-220 | [-1,96,36,36,36] | 192 |
| ReLU-221 | [-1,96,36,36,36] | 0 |
| UpsamplingBlock-222 | [-1,96,36,36,36] | 0 |
| Conv3d-223 | [-1,96,36,36,36] | 9,312 |
| BatchNorm3d-224 | [-1,96,36,36,36] | 192 |
| ReLU-225 | [-1,96,36,36,36] | 0 |
| Conv3d-226 | [-1,96,36,36,36] | 9,312 |
| BatchNorm3d-227 | [-1,96,36,36,36] | 192 |
| ReLU-228 | [-1,96,36,36,36] | 0 |
| TransitionBlock-229 | [-1,96,36,36,36] | 0 |
| Conv3d-230 | [-1,96,36,36,36] | 9,312 |
| BatchNorm3d-231 | [-1,96,36,36,36] | 192 |
| ReLU-232 | [-1,96,36,36,36] | 0 |
| Conv3d-233 | [-1,96,36,36,36] | 9,312 |
| BatchNorm3d-234 | [-1,96,36,36,36] | 192 |
| ReLU-235 | [-1,96,36,36,36] | 0 |
| TransitionBlock-236 | [-1,96,36,36,36] | 0 |
| Conv3d-237 | [-1,1,36,36,36] | 97 |
| Sigmoid-238 | [-1,1,36,36,36] | 0 |
