## Supplementary file for "DUnet: A deep learning guided protein-ligand binding pocket prediction"

**Tables**

**Table S1** Primers used in this study.

| Primers | Sequences (5′-3′) | Remarks |
| --- | --- | --- |
| 273f | GCGCACGGAAATCACTATCACGCGCCCAATGGCAG | Forward primer for generating mutation T273A. |
| 273r | CCAGACGGTCTTGTCGGCGTCCGCTTC | Reverse primer for generating mutation T273A. |
| 275f | GCGAATCACTATCACGCGCCCAATGGCAGCCT | Forward primer for mutation G275A. |
| 275r | GTGGGTCCAGACGGTCTTGTCGGCGTCC | Reverse primer for mutation G275A. |
| 289f | GCGGTCTACGAGAGCAAGTTCCGCAACTGGT | Forward primer for mutation H289A. |
| 289r | CATGGCACCCAGGCTGCCATTGGG | Reverse primer for mutation H289A |
| 290f | GCGTACGAGAGCAAGTTCCGCAACTGGT | Forward primer for mutation V290A. |
| 290r | ATGCATGGCACCCAGGCTGCCATTG | Reverse primer for mutation V290A. |
| 291f | GCGGAGAGCAAGTTCCGCAACTGGTCCGAGG | Forward primer for mutation Y291A. |
| 291r | GACATGCATGGCACCCAGGCTGCCA | Reverse primer for mutation Y291A. |
| 292f | GCGAGCAAGTTCCGCAACTGGTCCGAGGGT | Forward primer for mutation E292A. |
| 292r | GTAGACATGCATGGCACCCAGGCTGCCA | Reverse primer for mutation E292A. |
| 297f | GCGTGGTCCGAGGGTTACTCGGACTTC | Forward primer for mutation N297A. |
| 297r | GCGGAACTTGCTCTCGTAGACATGCA | Reverse primer for mutation N297A. |
| 298f | GCGTCCGAGGGTTACTCGGACTTCGAC | Forward primer for mutation W298A. |
| 298r | GTTGCGGAACTTGCTCTCGTAGACA | Reverse primer for mutation W298A. |
| 301f | GCGTACTCGGACTTCGACCGCGGAGCCTATGT | Forward primer for mutation N301A. |
| 301r | CTCGGACCAGTTGCGGAACTTGCTCTC | Reverse primer for mutation N301A. |

*The nucleotides generating mutation sites were underlined.

**Table S2** Success rate for k-fold validation.

|  | **Success rate** |
| --- | --- |
| **Fold-1** | **78%** |
| **Fold-2** | **84%** |
| **Fold-3** | **79%** |
| **Fold-4** | **80%** |
| **Fold-5** | **75%** |

* Success rate was defined as the distance between predicted binding area and actual ligand center below 4 Å.

**Figures**

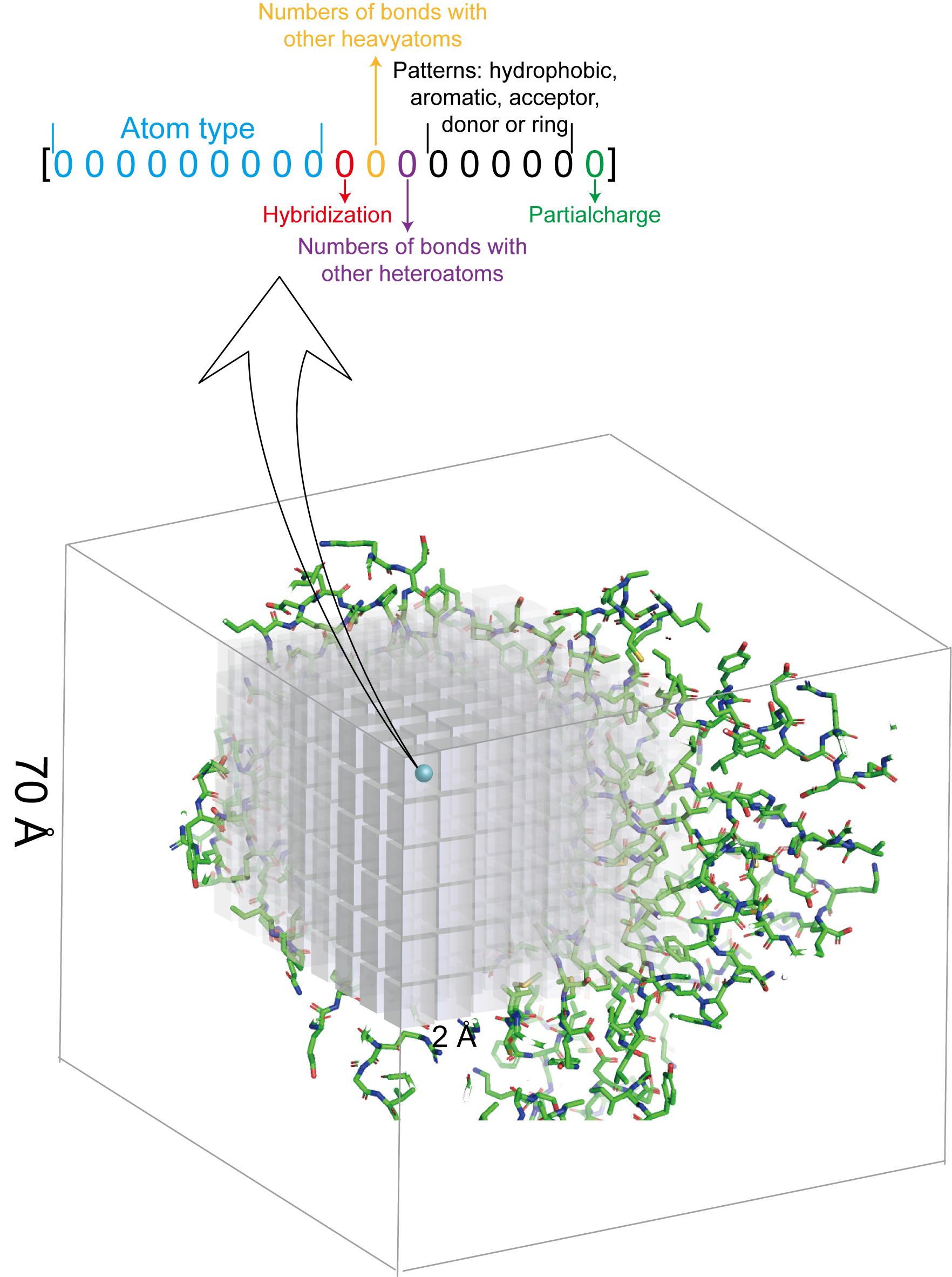

**Figure S1 Data representation**

Protein structure was treated as 3-dimentional image at a size of 70 Å^3^. The image was represented as voxels at a size of 2 Å^3^ to ensure each voxel can only contain one atom. Atomic features were described according to the physical characters.

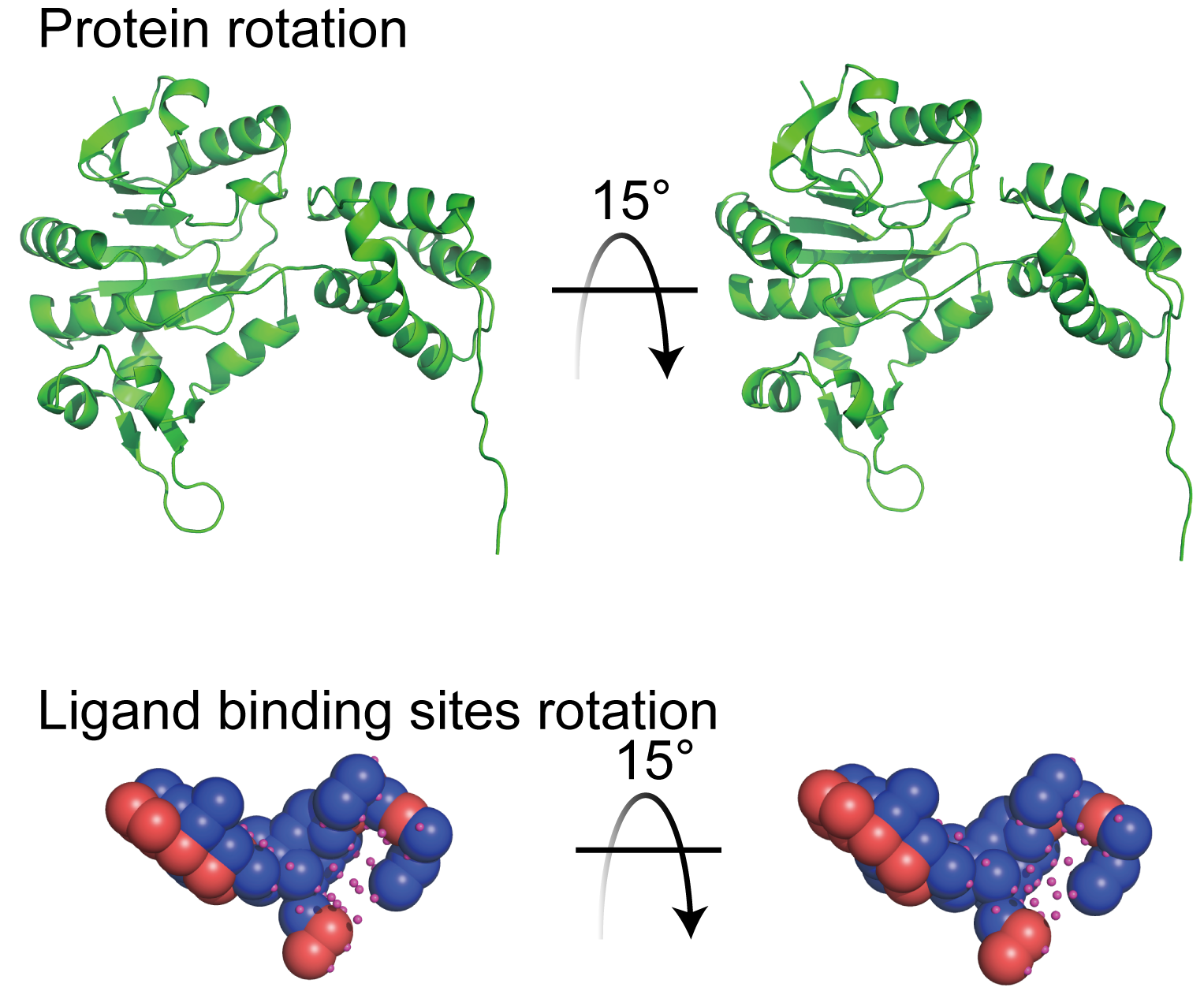

**Figure S2 Rotation of protein and ligand binding sites in scPDB_5020 dataset**

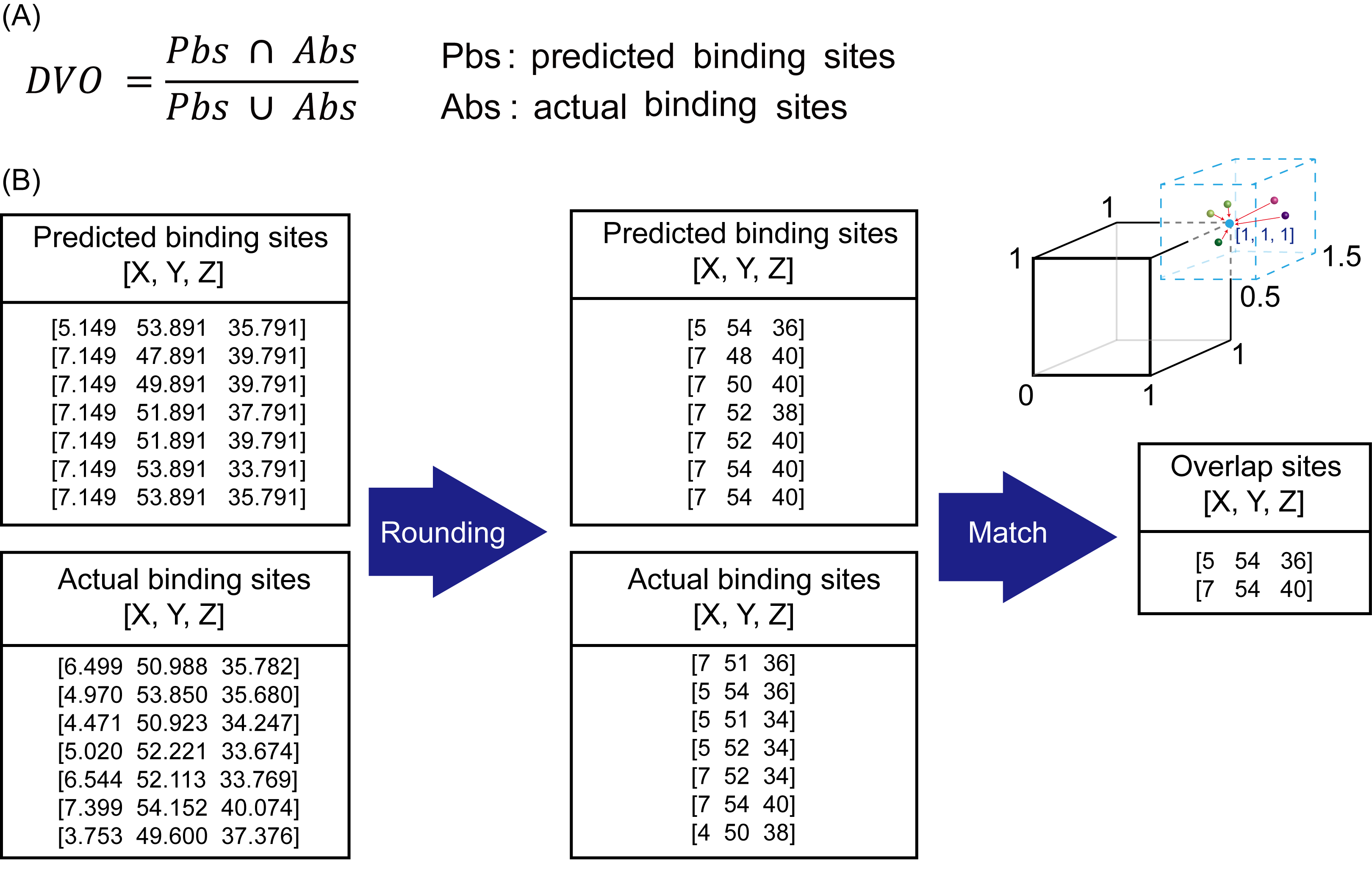

**Figure S3 Calculating the discretized volume overlap (DVO)**

(A) Mathematic representation of DVO calculation; (B) The rule for representing DVO.

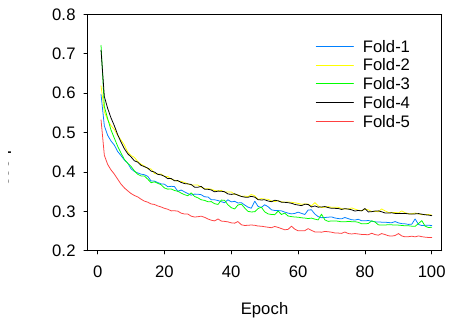

**Figure S4 Loss for k-fold validation**

scPDB_5020 subset was split into 5-fold for cross-validation, and 100 epochs were performed to train the model.
